# Innate Immune Correlates of Sickness-Like Depressive Symptoms in Breast Cancer Survivors

**DOI:** 10.64898/2026.10.12.758659

**Authors:** Iris Ka-Yi Chat, Steve W. Cole, Julienne E. Bower

## Abstract

**Background:** Research supports immune pathways of depression characterized by sickness-like symptoms (e.g., appetite changes, low energy, sleep disturbance, difficulty initiating activity, concentration difficulties), with limited evidence for negative affect symptoms (e.g., depressed mood, feeling of failure). However, most findings come from otherwise healthy individuals, limiting generalizability to medical populations. This study leveraged breast cancer as a disease context, given the lasting cancer- and treatment-related effects on immune function, to examine the hypothesis that proinflammatory gene expression relates more strongly to sickness-like than negative affect symptoms.

**Methods:** Data were drawn from 175 women with Stage 0-III breast cancer (M_age=_55.3 years) assessed at five time points from diagnosis through 18 months post-treatment. Depressive symptom dimensions were assessed using the Center for Epidemiologic Studies Depression Scale. Peripheral blood mononuclear cell gene expression was assessed using RNA sequencing-based transcriptional profiling. Primary analyses focused on proinflammatory gene expression. Exploratory analyses examined Type-I interferon gene expression and transcript-derived cell-type composition (myeloid-to-lymphoid and classical-to-non-classical monocyte ratios).

**Results:** Longitudinal mixed-effects models across repeated assessment revealed that greater sickness-like symptom severity was associated with elevated proinflammatory gene expression (p<.001), Type-I Interferon antiviral gene expression (p<.001), myeloid cell abundance (p<.001), and particularly non-classical monocytes (p=.002). By contrast, negative affect symptoms showed weaker, null, or opposite associations.

**Conclusions:** Sickness-like and negative affect symptoms show divergent associations with innate immune system activation and myeloid lineage bias. Results may inform more targeted interventions for both physical/energetic and psychological symptoms in breast cancer survivors, and may be relevant to other depression and medical contexts.

## Introduction

Depression is a major public health concern because of its high prevalence, recurrence, and role as a leading cause of disability worldwide (1). Depression is prominent among medically healthy populations and those living with major medical illnesses. For example, individuals with cancer experience higher rates of depression than matched primary care patients (2). Despite extensive research, depression remains difficult to treat, and a substantial proportion of individuals do not respond to available treatment (3). One reason may be that depression is highly heterogeneous and includes distinct symptom dimensions, which may have different biological underpinnings (4,5). However, depression is often studied and treated as a unitary disorder, which may obscure symptom-specific mechanisms and contribute to suboptimal treatment response. Identifying biological processes linked to specific depressive symptom dimensions may therefore help refine models of depression and inform more targeted approaches to treatment.

Emerging evidence suggests that elevated inflammation may be an underlying mechanism in a subset of individuals with depression (6,7). Inflammatory signaling can induce a coordinated set of behavioral changes during illness, often referred to as sickness behavior, including low effort-related motivation, low energy, appetite change, reduced psychomotor activity/speed, sleep disturbance, and concentration difficulties (8,9). These behaviors are thought to reflect an evolutionarily conserved response that promotes energy conservation and reallocates physiological resources toward immune defense and recovery. From this perspective, reduced motivation and increased somatic/vegetative symptoms (e.g., fatigue) may serve adaptative function by limiting energy expenditure and facilitating recovery (10). These sickness behaviors share considerable overlap with symptoms of depression, particularly vegetative and somatic symptoms such as fatigue, appetite change, sleep disruption, reduced activity/speed, and concentration difficulties. Importantly, prior research has consistently linked inflammation with these symptoms, including fatigue, appetite change, reduced motivation or effort-related behavior (8,9). Thus, inflammatory pathways may be especially relevant to sickness-like depressive symptom. In contrast, it remains less clear whether inflammatory biology is related to negative affect symptoms, such as sadness, emotional distress, and feelings of failure (9). Prior findings on inflammation and negative affect symptoms have been inconsistent (11–17). Yet, much of this work has examined overall depression severity (18,19) or individual symptoms in isolation (13,20), rather than directly comparing immune associations across theoretically distinct sickness-like and affective symptom dimensions. As such, the symptom specificity of immune-related processes remains unclear.

Clarifying this specificity also require immune assessments that extend beyond circulating inflammatory proteins. Most prior studies have focused on markers such as C-reactive protein (CRP) or interleukin-6 (7,18,21–23). These markers provide important information about inflammatory activity, but are sensitive to transient physiological changes (e.g., acute illness) and do not fully capture innate immune signaling pathways or regulatory processes, including those mediated by cellular mechanisms (e.g., circulating myeloid lineage cells involved in cell-mediated innate immune responses). Transcriptomic profiling of circulating leukocytes can capture broader profiles of proinflammatory signaling by evaluating expression of a coordinated set of proinflammatory genes. In addition, this approach can assess Type I interferon signaling, which remains less studied in relation to depressive symptoms (24–27) and is difficult to assess using circulating plasma proteins. Further, transcript-derived indicators of leukocyte subset composition may help distinguish whether symptom associations reflect inflammatory transcriptional activation within immune cells (e.g., transcript-derived classical monocytes) or shifts in the relative abundance of myeloid and lymphoid cell populations. Together, these transcriptomic approaches may provide a more comprehensive characterization of immune processes relevant to specific depressive symptom dimensions.

A second gap concerns the populations in which immune-related depressive symptom dimensions have been studied. Much of this literature has focused on medically healthy individuals with major depression (18,19,28). Although this approach reduces potential confounding by medical illness, it limits understanding of depressive symptoms in medically relevant contexts such as cancer survivorship. Depression is common among cancer patients and survivors (29,30), and depressive symptoms in these populations often include prominent sickness-like features (31–33). Meta-analyses have shown that depressive symptom severity is associated with protein markers of inflammation including tumor necrosis factor-α, interleukin-6, and CRP (34), with moderate-to-large effect size for these commonly studied markers. However, this work has largely focused on circulating inflammatory proteins and overall depressive symptom severity, and thus it remains unclear whether innate immune underpinnings differ across specific depressive symptom dimensions. Among the limited studies that have examined symptom dimensions in cancer populations, inflammatory markers such as interleukin-6 have been associated with somatic or vegetative symptoms but not affective symptoms (29,35,36). There is also evidence that specific somatic symptoms, especially cancer-related fatigue, are associated with elevated inflammatory activity (32,36–39). However, to our knowledge, no studies have directly examined whether innate immune transcriptional activity differentially associated with sickness-like versus affect-related depressive symptom profiles in cancer populations.

The current study was designed to examine innate immune correlates of sickness-related and negative affect-related depressive symptoms in a longitudinal study of women with early-stage breast cancer using transcriptional profiling of leukocytes. We have previously shown that fatigue was associated with elevated proinflammatory and Type I interferon gene expression in this sample (40). Here, we tested the hypothesis that greater proinflammatory gene expression would be associated with a more comprehensive constellation of sickness-like depressive symptoms, including fatigue/low energy, reduced appetite, difficulties getting going, reduced speech, sleep disturbance, and concentration difficulties. This symptom profile was compared with negative affect related symptoms, including sadness, depressed mood, fearfulness, emotional distress, inability to shake off the blues, and feelings of failure. To further characterize the innate immune processes linked to sickness-like depressive symptoms, we also conducted exploratory analyses examining Type I interferon-related gene expression and transcript-derived indices of myeloid lineage immune cell modulation. The transcript-derived indices provide estimates of leukocyte subset composition, including up-regulation of myeloid versus lymphoid lineage markers in general as well as alterations in the relative abundance or activity of classical versus non-classical monocyte markers. Together, these measures provide a more comprehensive approach for examining whether sickness-like depressive symptoms are associated not only with proinflammatory transcriptional activity, but also with broader antiviral and myeloid related immune processes.

## Methods and Materials

### Participants and procedures

Participants were recruited as part of the RISE longitudinal study of behavioral and biological correlates of cancer-related symptoms (39,41). The parent RISE Study enrolled 270 women from oncology clinics in Los Angeles between January 2013 and July 2015. Inclusion criteria were 1) diagnosis of Stage 0-IIIA breast cancer, 2) no prior initiation of adjuvant or neoadjuvant treatment (radiation therapy (RT), chemotherapy (CT), and/or endocrine therapy (ET)), and 3) English proficiency. The current analyses included participants who provided blood samples available for transcriptomic analyses and had available data on depressive symptoms and covariates; participants with comorbid autoimmune disease were excluded. This yielded an analytic sample of 175 women (M_age=_55.3 years). Sample characteristics are provided in Table 1.

**Table 1.**
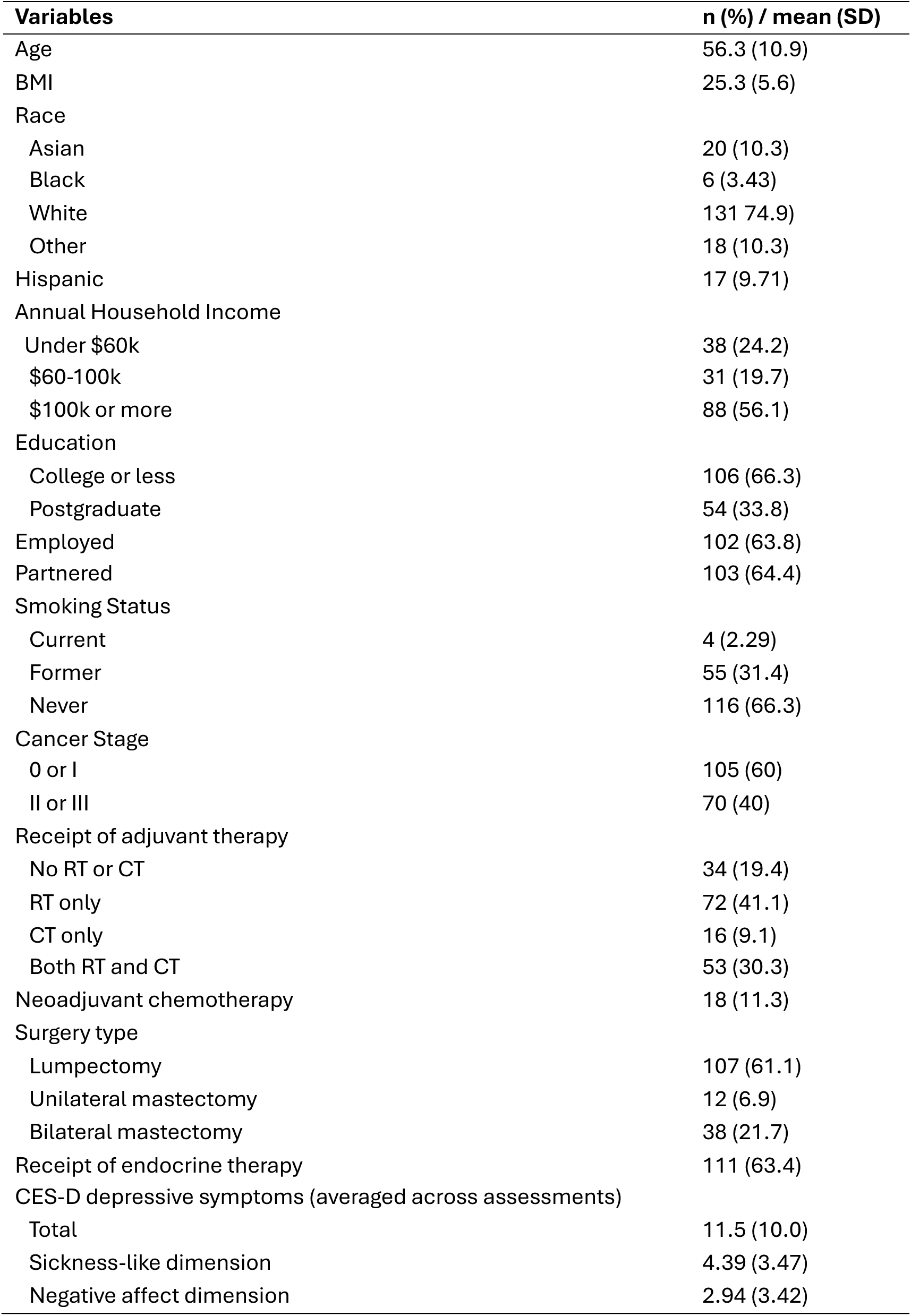
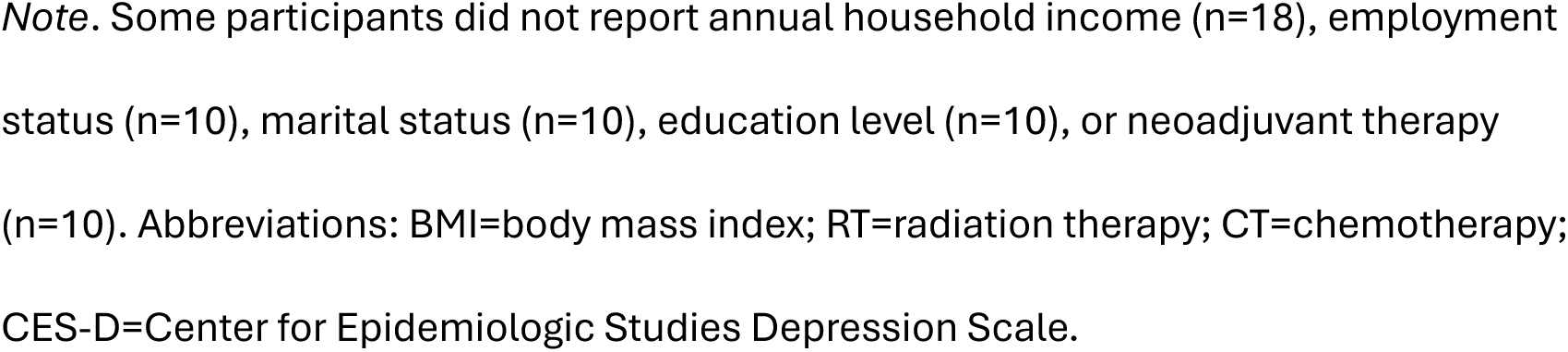
Sample characteristics (N=175).

| **Variables** | **n (%) / mean (SD)** |
| --- | --- |
| Age | 56.3 (10.9) |
| BMI | 25.3 (5.6) |
| Race |  |
| Asian | 20 (10.3) |
| Black | 6 (3.43) |
| White | 131 (74.9) |
| Other | 18 (10.3) |
| Hispanic | 17 (9.71) |
| Annual Household Income |  |
| Under \$60k | 38 (24.2) |
| \$60-100k | 31 (19.7) |
| \$100k or more | 88 (56.1) |
| Education |  |
| College or less | 106 (66.3) |
| Postgraduate | 54 (33.8) |
| Employed | 102 (63.8) |
| Partnered | 103 (64.4) |
| Smoking Status |  |
| Current | 4 (2.29) |
| Former | 55 (31.4) |
| Never | 116 (66.3) |
| Cancer Stage |  |
| 0 or I | 105 (60) |
| II or III | 70 (40) |
| Receipt of adjuvant therapy |  |
| No RT or CT | 34 (19.4) |
| RT only | 72 (41.1) |
| CT only | 16 (9.1) |
| Both RT and CT | 53 (30.3) |
| Neoadjuvant chemotherapy | 18 (11.3) |
| Surgery type |  |
| Lumpectomy | 107 (61.1) |
| Unilateral mastectomy | 12 (6.9) |
| Bilateral mastectomy | 38 (21.7) |
| Receipt of endocrine therapy | 111 (63.4) |
| CES-D depressive symptoms (averaged across assessments) |  |
| Total | 11.5 (10.0) |
| Sickness-like dimension | 4.39 (3.47) |
| Negative affect dimension | 2.94 (3.42) |

*Note.* Some participants did not report annual household income (n=18), employment status (n=10), marital status (n=10), education level (n=10), or neoadjuvant therapy (n=10). Abbreviations: BMI=body mass index; RT=radiation therapy; CT=chemotherapy; CES-D=Center for Epidemiologic Studies Depression Scale.

Participants completed baseline assessments after breast cancer diagnosis and before the start of adjuvant or neoadjuvant treatment, including RT, CT, and/or ET. Additional assessments were conducted after completion of RT and/or CT among participants who received those treatments (post-treatment) and at 6-, 12-, and 18-month post-treatment follow-ups, capturing acute- and longer-term changes associated with cancer treatments and recovery. At each visit, participants completed self-report questionnaires and provided blood samples via venipuncture for immune assessment.

All study procedures were approved a priori by the Institutional Review Boards at both UCLA and Cedar Sinai Medical Center, and written informed consent was obtained from all participants.

## Measures

### Demographic Characteristics, Anthropometrics, and Disease and Treatment-Related Information

Demographic and anthropometric information was obtained at the baseline visit. This includes age, race, ethnicity, body mass index, and smoking status. Disease and treatment-related information, including cancer stage, type of surgery (lumpectomy, unilateral mastectomy, bilateral mastectomy), and type of adjuvant therapy received (radiation therapy, chemotherapy, and/or endocrine therapy), was acquired from medical record abstraction.

### Depressive Symptom Dimensions

The Center for Epidemiologic Studies Depression Scale (CES-D) (42) is a 20-item self-report measure assessing depressive symptoms experienced during the past week. Although developed as a screening instrument rather than a diagnostic assessment, the CES-D has demonstrated good accuracy among adults, including patients with chronic medical conditions (43,44). Items are rated on a 4-point (0–3) scale, with total scores ranging from 0 to 60.

***Sickness-like and negative affect symptom dimensions*.**

To distinguish depressive symptoms most closely aligned with sickness behavior from affective symptoms of depression, two CES-D symptom dimensions were derived using a conceptually and empirically informed approach. Sickness-like symptoms were identified based on prior sickness behavior theory and inflammation-associated behavioral phenotypes (9,45), and were further evaluated using confirmatory factor analysis in the present sample. The resulting six-item sickness-like depressive symptom score was computed from CES-D items including fatigue/low energy, reduced appetite, reduced speech, difficulty getting going, sleep disturbance, and concentration difficulties (See Supplementary Table S1 for specific items)(8,9). A six-item negative affect score was computed from CES-D items reflecting sadness, depressed mood, fearfulness, distress, inability to shake off the blues, and feeling of failure (Supplementary Table S1)(46). For both scales, higher scores indicate greater symptom severity.

These dimensions overlap with, but are not identical to, traditional CES-D Somatic/Vegetative and Depressed Affect subscales (46)(see Supplementary Table S1 for specific items and comparison). In the present sample, confirmatory factor analysis indicated good fit for the proposed sickness-like and negative affect symptom structure (CFI=.975 vs .944, TLI=.969 vs .933; RMSEA=.046 vs.066). For descriptive comparison, we also examined the traditional CES-D Somatic/Vegetative and Depressed Affect subscale structure, which yielded CFI = .944, TLI = .933, and RMSEA = .066.

### Gene Expression Immune Indices

Peripheral blood mononuclear cells (PBMCs) were isolated from whole blood sample collected by venipuncture, and total RNA was extracted from PBMCs using Qiagen RNeasy kits. Genome-wide mRNA profiling was performed using QuantSeq 3′ FWD cDNA library synthesis (Lexogen) and sequencing on an Illumina HiSeq 4000 instrument at the UCLA Neuroscience Genomics Core, all following manufacturer protocols. Sequencing reads were aligned to the GRCh38 human reference transcriptome using STAR aligner, with transcript abundance quantified as log2-transformed counts per million mapped reads. Quality control procedures excluded samples with fewer than 5 million total sequencing reads, mapping rates below 70%, or low transcriptome profile concordance with other samples (median correlation <.80). In addition, three of the a priori-targeted gene transcripts with zero variability were excluded from analyses. Complete details on genome-wide transcriptional profiling methods are provided in our previous report (47).

Analyses focused on two predefined gene composites reflecting inflammatory and antiviral transcriptional activity. These composites were selected given that they represent functionally distinct innate regulatory processes implication in chronic disease risk and have been used in prior research to capture broad variation in leukocyte proinflammatory and Type I interferon activity (48). The proinflammatory composite included 19 genes (*CXCL8, FOS, FOSB, FOSL1, FOSL2, IL1A, IL1B, IL6, JUN, JUNB*, *JUND, NFKB1, NFKB2, PTGS1, PTGS2, REL, RELA, RELB, and TNF*) (48)), and the antiviral composite included 31 genes (*GBP1, IFI16, IFI27, IFI27L1, IFI27L2, IFI30, IFI35, IFI44, IFI44L, IFI6, IFIH1, IFIT1, IFIT1B, IFIT2, IFIT3, IFIT5, IFITM1, IFITM2, IFITM3, IFITM4P, IGLL1, IRF2, IRF7, IRF8, JCHAIN, MX1, MX2, OAS1, OAS2, OAS3, and OASL*)(48).

In addition, transcript-based markers were used to estimate leukocyte subset composition. Cell subset markers were provided from RNA-sequencing data and reflected relative expression of canonical leukocyte markers. A monocyte subset bias ratio, computed as *CD14-FCGR3A* (CD16), indexes classical monocyte relative to non-classical monocyte abundance. A broader myeloid-to-lymphoid ratio, computed as the average expression of myeloid markers (*CD14* and *FCGR3A*) minus the average expression of lymphoid markers (*CD3D*, CD19, *NCAM1*), indexes relative abundance of monocyte markers relative to T-cell, B-cell, and natural killer cell abundance. Higher values indicate greater relative expression of the first cell class (i.e., classical monocyte and myeloid lineage, respectively) in each contrast.

### Statistical Analyses

Linear mixed-effects model analyses were conducted to examine associations between innate immune activation and depressive symptoms within each dimension, and whether they differed across sickness-like vs. affective symptom dimensions. Models used all available repeated symptom observations across up to five assessment visits. The longitudinal symptom data in this study included 1,086 symptom-dimension observations from 544 assessment visits among 175 individuals. The immune indices x symptom type interaction term provided the formal test of whether the associations between immune measures and symptom severity differed significantly between sickness-like and negative affect symptoms. Primary analyses focused on proinflammatory gene expression scores. Exploratory analyses repeated these models using Type I interferon gene expression, myeloid-to-lymphoid ratios, and classical-to-non-classical monocyte ratios.

All models adjusted for factors that may be associated with the immune and clinical variables as covariates (39,49), including age, race, ethnicity, body mass index, smoking status, cancer stage (0/I vs. II/III), surgery type, and cancer treatment (chemotherapy, radiation, endocrine therapy). Assessment time point was also included to account for longitudinal assessments across the study period. Models involving proinflammatory and Type I interferon gene expression additionally adjusted for transcript-derived indicators of leukocyte subset abundance (*CD14, CD3D, CD4, CD8A, CD19,* CD56/*NCAM1*, CD16/*FCGR3A*) to isolate effects of transcriptional activity independent of differences in transcript-level leukocyte composition. Models included a random participant intercept to account for repeated observations within participants. Gene was included as a fixed effect to account for systematic differences across indicator genes. Analyses were conducted using R software.

## Results

### Characteristics of Depressive Symptom Dimensions Across Assessment Time Points

To characterize sickness-like and negative affective depressive symptom dimensions across the study period, we examined symptom severity across assessment time points over the first 18 months of breast cancer survivorship. Mean sickness-like depressive symptoms were consistently higher than negative affect symptoms across time points, F(1, 895)=9.32, p=.002, that is, at baseline (4.75 vs. 3.49), post-treatment (4.67 vs. 2.73), 6-month follow-up (4.74 vs. 3.33), 12-month follow-up (4.49 vs. 2.85), and 18-month follow-up (4.23 vs. 2.75; Figure 1). Symptom dimension severity did not significantly vary across time, F(4, 895), 1.41, p=.227), and the difference between sickness-like and negative affect symptoms did not significantly change over time (symptom dimension x time interaction: F(4, 895)=.59, p=.673. Among covariates, younger age was associated with greater overall symptom severity, F(1,895)=6.89, p=.009. In dimension-specific models, younger age was associated with greater sickness-like symptom severity, F(1, 362)=6.93, p=.009, and greater negative affect symptom severity, F(1, 360)=5.42, p=.021. These associations did not significantly differ by symptom dimension, F(1, 894)=.008, *p*=.928. Other demographic and clinical covariates were not significantly associated with overall symptom severity, F’s≤1.96, p’s≥.122. They were also not significantly associated with symptom severity in dimension-specific models, sickness-like symptoms: F’s≤3.43, p’s≥.066; negative affect symptoms: F’s≤2.34, p’s≥.076.

**Figure 1.**
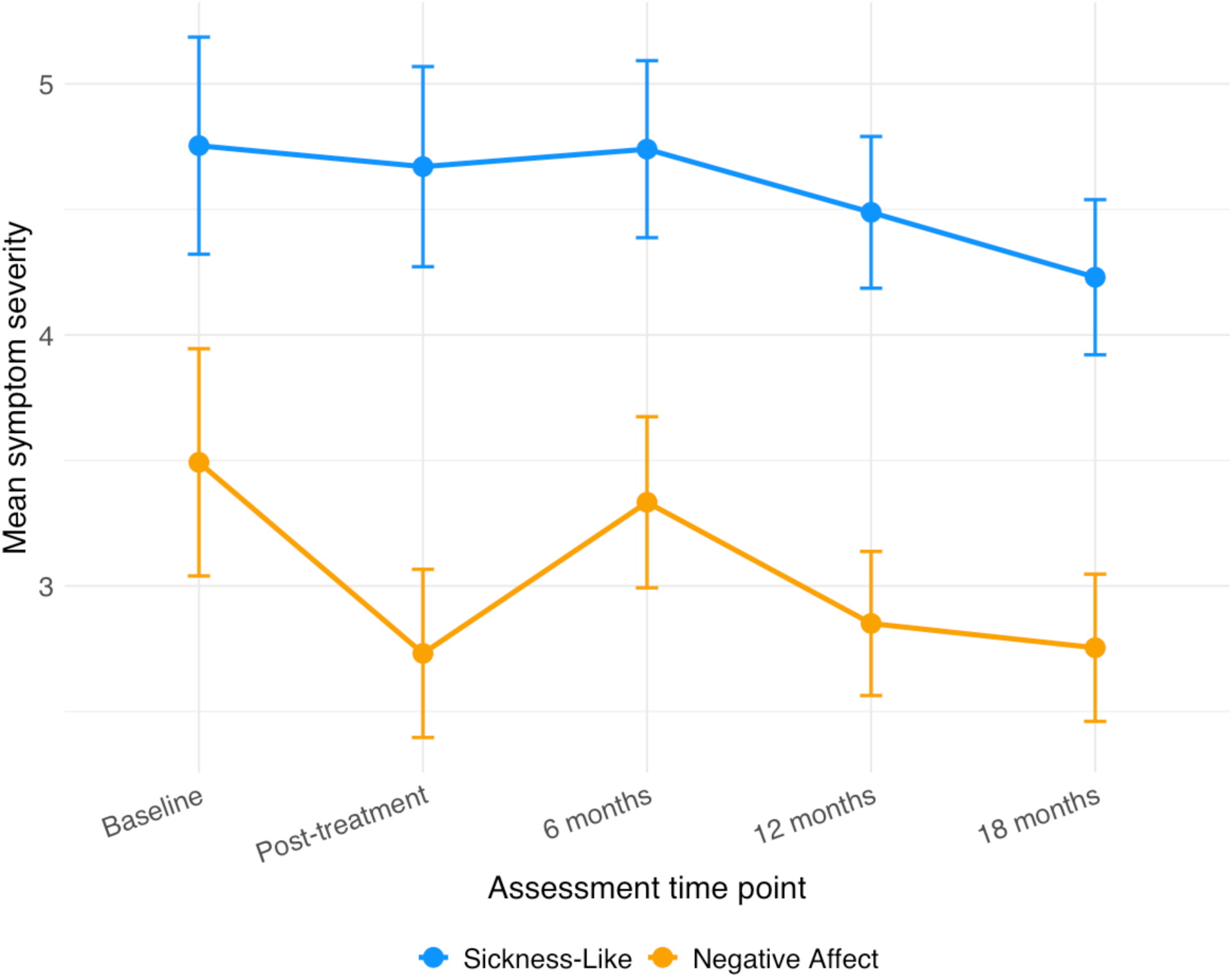
Sickness-like and negative affect depressive symptoms across assessment visits. *Note*. Points represent mean symptom severity at each assessment visit. Error bars represent standard errors. Sickness-like symptoms were consistently higher than negative affect symptoms across the study period. Symptom severity did not significantly vary over time and the symptom dimension x time interaction was not significant.

### Innate Immune Regulation and Depressive Symptom Dimensions Across Assessment Time Points

#### Proinffammatory Gene Expression

We next examined the primary study hypothesis that pro-inflammatory gene expression showed differential associations with sickness-like versus negative affective symptoms of depression. As hypothesized, greater proinflammatory gene expression was associated with greater sickness-like depressive symptoms (*b*=.10, *SE*=.02, *p*<.001). In contrast, the association with negative affect symptoms was less than half that magnitude and in the opposite direction (*b*=-.04, *SE*=.02, *p*=.055) (Figure 2). This differential associations across symptom dimensions were confirmed by a significant proinflammatory gene expression x symptom dimension interaction effect, *F*(1, 20297)=22.45, *p*<.001. Similar results emerged after additionally adjusting for transcript-derived indicators of leukocyte subset abundance (sickness-like symptoms: *b*=.08, *SE*=.02, *p*=.001; negative affect symptoms: *b*=-.07, *SE*=.02, *p*=.003), suggesting that differential leukocyte subset abundance was not the sole mechanism for these associations.

**Figure 2.**
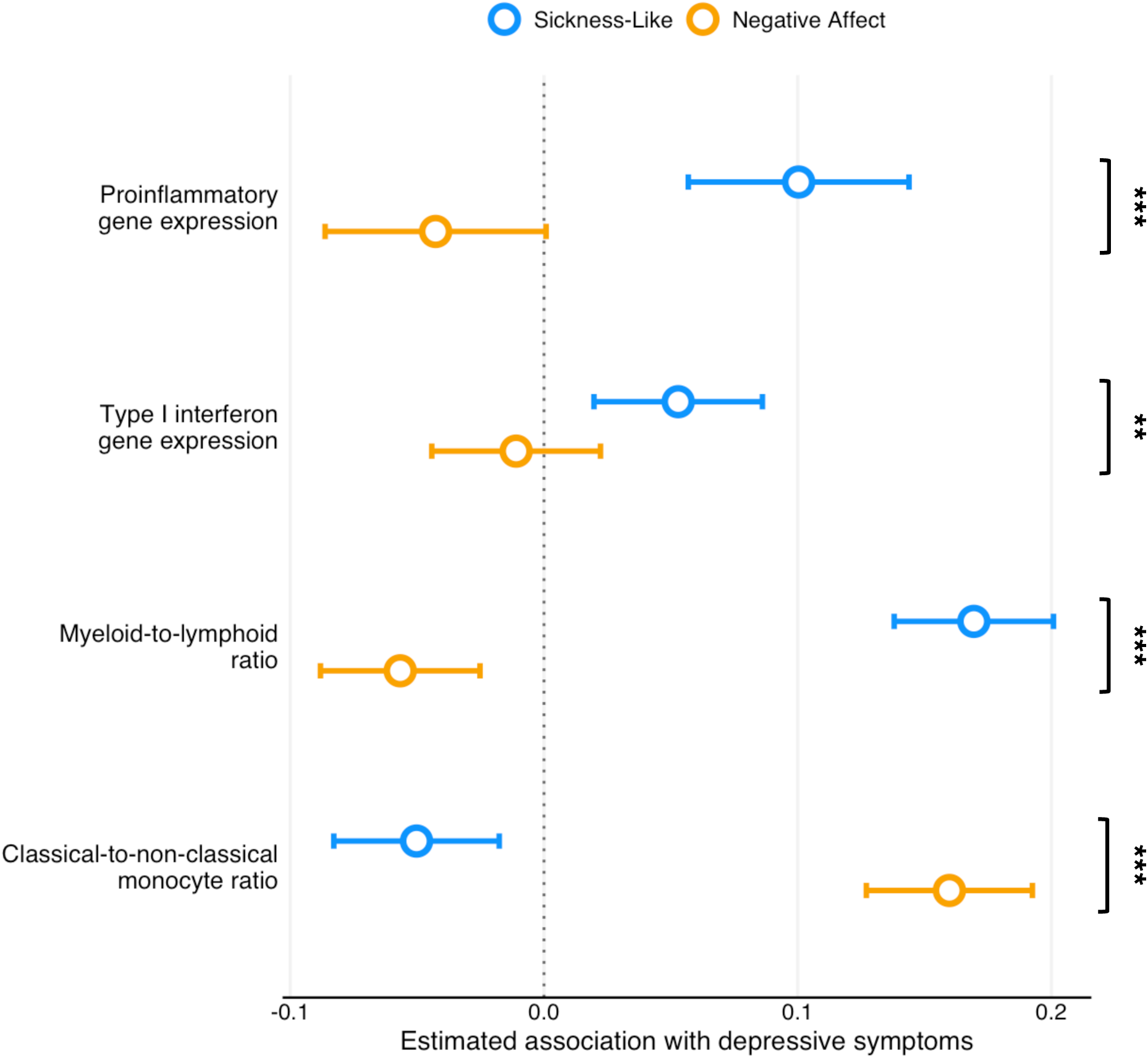
Associations between Immune Indices and Depressive Symptom Dimensions. *Note*. Points represent estimated simple slopes for the association between each immune index (proinflammatory gene expression, Type I interferon gene expression, myeloid-to-lymphoid lineage ratio, classical-to-non-classical monocyte ratio,) and depressive symptom severity across dimensions (sickness-like vs. negative affect symptoms). Error bars represent 95% confidence intervals. Asterisks indicate significance of the immune index x symptom dimension interaction, reflecting that associations between each immune index and symptom severity differed between sickness-like and negative affect symptom dimensions (\*\**p* <.01, \*\*\**p* <.001).

#### Type I Interferon Gene Expression

To determine whether the antiviral dimension of innate immunity might also show differential associations with sickness-like versus affective symptoms of depression, analyses were conducted with Type I interferon gene expression as a predictor of depressive symptom dimensions. Consistent with findings for proinflammatory gene expression, greater Type I interferon gene expression was associated with more severe sickness-like depressive symptoms (*b*=.053, *SE*=.017, *p*=.002). Again, associations with negative affect symptoms were minimal, in the opposite direction, and not significant (*b*=-.01, *SE*=.02, *p*=.517) (Figure 2). This differential pattern was confirmed by a significant Type I interferon gene expression x symptom dimension interaction effect, *F*(1, 33233)=7.36, *p*=.007. Similar results emerged after additionally adjusting for transcript-derived indicators of leukocyte subset abundance (sickness-like symptoms: *b*=.03, *SE*=.02, *p*=.05; negative affect symptoms: *b*=-.03, *SE*=.02, *p*=.07), suggesting that differential leukocyte subset abundance was not the sole mechanism underlying these associations.

#### Myeloid Cell Subsets

To determine whether changes in the myeloid lineage cellular composition of circulating blood might contribute to the differential associations of innate immune response gene expression with sickness-like versus negative affective symptoms of depression, we conducted additional analyses using a measure of myeloid lineage bias in the circulating PBMC pool (i.e., the ratio of monocyte markers to lymphocyte markers). Greater myeloid-to-lymphoid lineage ratio was associated with more severe sickness-like depressive symptoms (*b*=.19, *SE*=.02, *p*<.001), whereas the association with negative affect symptoms was less than half the magnitude and in the opposite direction (*b*=-.06, *SE*=.02, *p*=.001) (Figure 2). These differential associations of sickness-like versus affective symptoms of depression were confirmed by a significant lineage ratio x symptom dimension interaction, *F*(1, 53715)=140.05, *p*<.001.

Finally, we asked whether the overall myeloid lineage association with sickness-like symptoms might reflect a change in cell differentiation status within the myeloid lineage cell pool – specifically, a change in the relative prevalence of classical vs. non-classical monocytes. A relative abundance of non-classical monocytes (i.e., lower classical-to-non-classical monocyte ratio) was associated with significantly more severe sickness-like depressive symptoms (*b*=-.05, *SE*=.02, *p*=.003), but significantly less negative affect symptoms (*b*=.16, *SE*=.02, *p*<.001) (Figure 2). Again this differential association with sickness-related versus negative affective symptoms of depression was confirmed by a significant lineage ratio x symptom dimension interaction, *F*(1, 53715)=125.93, *p*<.001.

## Discussion

This longitudinal study demonstrated that activation of the innate immune system was distinctively associated with sickness-like depressive symptoms among women with early-stage breast cancer. Sickness-like depressive symptoms were linked to multiple aspects of innate immune system activation, including the expression of pro-inflammatory genes, as well as the expression of Type I interferon response genes, the relative abundance of myeloid lineage immune cells relative to lymphoid lineage cells, and a selective up-regulation of non-classical monocytes within the circulating monocyte pool. These effects were relatively specific to sickness-like symptoms, including appetite changes, low energy, difficulty with effort-based behavior, reduced speech, sleep disturbance, and concentration difficulties, as these same measures of innate immune system activation showed strikingly different associations with negative affective symptoms of depression (generally showing either no association or associations in the opposite direction). These results suggest a specific link between innate immune system activation and sickness-related depressive symptoms in the context of breast cancer treatment and recovery, and they highlight the importance of distinguishing depressive symptom dimensions in research on immune-related depression.

These findings extend prior studies in cancer patients and cancer survivors linking inflammation with somatic or vegetative symptoms, including fatigue and other individual sickness-like behavioral symptoms (29,32,36–39,50). More specifically, the limited studies that have examined depressive symptom dimensions in cancer populations suggest that circulating inflammatory markers, such as interleukin-6, are more consistently associated with somatic/vegetative symptom severity than with affective symptoms (29,35). The present study builds on this work in several ways. First, we examined an *a priori*, empirically supported sickness-like depressive symptom dimension grounded in sickness behavior theory (45) and prior integrative research (8,9). This approach is useful because the sickness-like symptom dimension was selected to reflect a coordinated behavioral phenotype more closely aligned with immune-related sickness behavior as described in sickness behavior theories (9,45). Second, we expanded immune assessment beyond circulating inflammatory proteins by examining leukocyte transcriptional profiles, including proinflammatory gene expression, Type I interferon gene expression, and transcript-derived indicators of myeloid lineage and monocyte subset composition. Finally, the longitudinal design and relatively large sample of breast cancer survivors allowed for examining these associations across periods of diagnosis through treatment and recovery.

The contrasting associations observed across sickness-like and negative affect symptoms support the view that inflammatory activity may be differentially linked to specific depressive symptom dimensions, rather than depression uniformly. Although limited prior studies suggest that negative affect symptoms may relate to inflammatory processes (8,14–16), other studies have not found consistent associations (9,11–13). In the current sample, negative affect symptoms were not significantly associated with proinflammatory or Type I interferon gene expression. Moreover, associations between negative affect symptoms and transcript-derived cell composition indices were in the opposite direction from those observed for sickness-like symptoms. These findings are consistent with prior longitudinal work by our group showing that circulating inflammatory proteins were positively associated with general and physical fatigue, not associated with mental fatigue, and negatively associated with emotional fatigue, suggesting that greater inflammatory activity may be more closely linked to physical and sickness-like symptom dimensions than to emotional distress (51). Together, this broader pattern underscores the importance of examining immune-symptom associations longitudinally and dimensionally, rather than treating depression as a unitary clinical endpoint or relying solely on diagnostic group comparisons. These findings do not rule out a role for immune processes in negative affect, but they suggest that negative affect symptoms may show a distinct immune profile from sickness-like depressive symptoms, at least in the context of breast cancer. Future research should clarify whether immune processes contribute to negative affective symptoms directly, indirectly through related mechanisms, or only in specific clinical or biological contexts. Notably, these associations were observed in a sample that included both clinical and subthreshold depressive symptoms, consistent with a dimensional view in which immune-sickness symptom links may be evident even outside the context of clinically significant depression.

In addition to testing symptom specificity, this study extends prior work by examining multiple transcript-derived indicators of innate immune activity. Prior studies of cancer and depressive symptoms have often relied on circulating inflammatory proteins, although emerging research has examined immune gene expression and other molecular immune processes in relation to fatigue and related depressive symptoms (40,52,53). Here, sickness-like depressive symptoms were associated not only with proinflammatory gene expression, but also with Type I interferon gene expression and transcript-derived indices of myeloid lineage activity. This pattern suggests that sickness-like depressive symptoms may be linked to a broader, coordinated innate immune phenotype involving proinflammatory, antiviral, and myeloid regulatory processes (54), rather than proinflammatory signaling alone. Of note, the associations with myeloid-to-lymphoid and monocyte subset indices further suggest that this profile may involve not only immune transcriptional activity, but also inferred differences in leukocyte subset composition. Together, these findings highlight the value of transcriptomic approaches for capturing partially overlapping aspects of immune biology relevant to sickness-like depressive symptoms, including inflammatory signaling, antiviral signaling, and inferred leukocyte subset composition.

Several limitations should be considered. First, participants are predominantly non-Hispanic White, which may limit the generalizability of the present findings to other racial and ethnic groups. This limitation may be particularly important given evidence of racial and ethnic differences in both inflammatory processes and depression (55,56). Second, treatment modalities may differentially influence immune function (57), and the present study was underpowered to comprehensively examine treatment-specific effects beyond including treatment exposure as a covariate. Although this approach helped reduce potential confounding, the clinical implications of different treatment exposures remain unclear. Third, depressive symptoms in this sample were largely subthreshold, which may limit generalizability to individuals meeting clinical thresholds for depression. Moreover, CES-D-derived symptom dimensions were based on self-report, and the negative affect dimension did not capture the full range of symptoms. Relatively low negative affect scores may also have limited sensitivity to detect associations with immune measures. Although the proposed symptom were theoretically informed and support by CFA, their measurement invariance, reliability, and convergent and discriminant validity were not formally evaluated. Thus, these dimensions should be considered preliminary and require further psychometric validation. Finally, the observational nature of the study precludes causality claims about the direction of associations between immune alterations and depressive symptoms.

## Conclusions

These findings identified differential associations between innate immune system activation and sickness-like versus negative affect depressive symptoms in the context of treatment and recovery among breast cancer survivors. They highlight the importance of examining depressive symptom dimensions in studies of depression syndrome-level biology and suggest that medically relevant immune contexts may provide useful models for understanding symptom-specific immune processes. Future research should examine these associations across populations with other medical conditions and clarify which aspects of survivorship contribute to immune-related depressive symptom vulnerability.

## Acknowledgments

This work was funded by National Institutes of Health/National Cancer Institute R01CA160427 (PI: JEB), T32MH015750 (IKC), and the Breast Cancer Research Foundation. During the preparation of this work, the authors used ChatGPT to assist with language editing and refinement of human-written manuscript text. After using this tool, the authors carefully reviewed and edited the content as needed, and the authors take full responsibility for the content of the article.

## Disclosure of Biomedical Financial Interests and Potential Conflicts of Interest

The authors reported no biomedical financial interests or potential conflicts of interest.

## Supplementary Material

**Table S1.** Center for Epidemiologic Studies Depression Scale (CES-D) items included in sickness-like, negative affect, somatic, and depressed affect symptom dimensions.

| No. | CES-D Item | Sickness-Like | Negative Affect | Somatic / Vegetative | Depressed Affect |
| --- | --- | --- | --- | --- | --- |
| 1 | I was bothered by things that usually do not bother me |  | x | x |  |
| 2 | My appetite was poor | x |  | x |  |
| 3 | I could not shake off the blues even with help from my family or friends |  | x |  | x |
| 4 | I had trouble keeping my mind on what I was doing | x |  | x |  |
| 5 | I felt depressed |  | x |  | x |
| 6 | Everything I did was an effort | x |  | x |  |
| 7 | I thought my life had been a failure |  | x |  | x |
| 8 | I felt fearful |  | x |  | x |
| 9 | My sleep was restless | x |  | x |  |
| 10 | I talked less than usual | x |  | x |  |
| 11 | I felt lonely |  |  |  | x |
| 12 | I had crying spells |  |  |  | x |
| 13 | I felt sad |  | x |  | x |
| 14 | I could not get going | x |  | x |  |

*Note.* Sickness-like and negative affect symptom dimensions were based on prior work
distinguishing inflammation-associated sickness-like symptoms from negative affective symptoms
(1). Relative to the traditional CES-D Somatic/Vegetative and Depressed Affect subscales (2), the sickness-like dimension excludes Item 1 ("I was bothered by things that usually do not bother me"), which was instead included in the negative affect symptom dimension because it aligns with emotional distress rather than prototypical sickness behavior. The negative affect dimension excludes Items 14 ("I felt lonely") and 17 ("I had crying spells") to focus more specifically on core negative emotional symptoms. Confirmatory factor analyses in the present sample supported the proposed sickness-like and negative affect symptom structure (CFI = .975, TLI = .969, RMSEA = .046, SRMR = .048), which showed better fit than the traditional Somatic and Depressed Affect structure (CFI = .944, TLI = .933, RMSEA = .066, SRMR = .049). Abbreviation: CES-D=Center for Epidemiologic Studies Depression Scale.

